# Neural Taskonomy: Inferring the Similarity of Task-Derived Representations from Brain Activity

**DOI:** 10.1101/708016

**Authors:** Aria Y. Wang, Leila Wehbe, Michael J. Tarr

## Abstract

Convolutional neural networks (CNNs) trained for object recognition have been widely used to account for visually-driven neural responses in both the human and primate brains. However, because of the generality and complexity of the task of object classification, it is often difficult to make precise inferences about neural information processing using CNN representations from object classification despite the fact that these representations are effective for predicting brain activity. To better understand underlying the nature of the visual features encoded in different brain regions of the human brain, we predicted brain responses to images using fine-grained representations drawn from 19 specific computer vision tasks. Individual encoding models for each task were constructed and then applied to BOLD5000—a large-scale dataset comprised of fMRI scans collected while observers viewed over 5000 naturalistic scene and object images. Because different encoding models predict activity in different brain regions, we were able to associate specific vision tasks with each region. For example, within scene-selective brain regions, features from 3D tasks such as 3D keypoints and 3D edges explain greater variance as compared to 2D tasks—a pattern that replicates across the whole brain. Using results across all 19 task representations, we constructed a “task graph” based on the spatial layout of well-predicted brain areas from each task. We then compared the brain-derived task structure with the task structure derived from transfer learning accuracy in order to assess the degree of shared information between the two task spaces. These computationally-driven results—arising out of state-of-the-art computer vision methods—begin to reveal the task-specific architecture of the human visual system.

## 1 Introduction

Scene understanding requires the integration of space perception, visual object recognition, and the extraction of semantic meaning. The human brain’s solution to this challenge has been elucidated in recent years by the identification of scene-selective brain areas via comparisons between images of places and common objects^[1]^. This basic contrast has been refined across a wide variety of image manipulations that have provided evidence for the neural coding of scene-relevant properties such as relative openness^[2–4]^, the distance of scenes to the viewer^[2,3,5]^, the spatial layout of 3D space^[6–8]^ and navigational affordances^[9]^. Recently, to help explain such findings, Lescroart and Gallant^[10]^ developed an encoding model using a feature space that parametrizes 3D scene structures along the distance and orientation dimensions and provides a computational framework to account for human scene processing. Intriguingly, Lescroart and Gallant^[10]^ were able to identify distance and openness within scenes as the dimensions that best account for neural responses in scene-selective brain areas. At a higher, semantic, level, Stansbury et al.^[11]^ found that neural responses in scene-selective brain areas can be predicted using scene categories that were learned from object co-occurrence statistics. Such findings demonstrate that human scene-selective areas represent both visual and semantic scene features. At the same time, there is still no robust model of how these different kinds of information are integrated both within and across brain regions.

Encoding models are widely used in understanding feedforward information processing in human perception, including scene perception. Encoding models are predictive models of brain activity that are able to generalize and predict brain responses to novel stimuli. Researchers have also used encoding models to infer which dimensions are critical for prediction by comparing the weights learned by the model. One of the successes of encoding models lies in predicting low-to mid-level visual cortex responses in humans and primates using features that were learned via a convolutional neural network trained on object recognition^[12–15]^. Most interestingly, these studies demonstrate a correspondence between human neural representation and learned representations within CNN models along the perceptual hierarchy: early layers tend to predict early visual processing regions, whereas later layers tend to predict later visual processing regions. Similarly, researchers have found that network representations from other task-driven networks, including networks trained on speech- or music-related tasks, are able to explain neural responses in human auditory pathways^[16]^. Such successes are not mere coincidences but rather indications of how fundamental task-driven representations are to both task training and to information processing in the brain.

Despite these advances, CNN features themselves are notoriously difficult to interpret. First, activations from the convolutional layers lie in extremely high-dimensional space and it is difficult to interpret what each feature dimension signifies. Second, features from a CNN tailored for a particular visual task can represent any image information that is relevant to that task. As a consequence of these two issues, the feature representations learned by the network are not necessarily informative with respect to the nature of visual processing in the brain despite their good performance in predicting brain activity.

To better understand the specificity of the information represented in the human visual processing pathways, we adopted a different approach. Instead of choosing a generic object-classification CNN as a source of visual features, we built encoding models with individual feature spaces obtained from different task-specific networks. These tasks included mid-level features such as surface normal estimation, edge detection, scene classification, *et. cetra*. In any task-driven network, the feature space learned to accomplish the task at hand should only represent information from input images that is task-relevant. Therefore we can use the predictive regions from each of the models to identify the brain regions where specific task-relevant information is localized. Independently, Dwivedi and Roig^[17]^ has shown that representation similarity analysis (RSA) performed between task representations and brain representations can differentiate scene-selective regions of interest (ROIs) by their preferred task. For example, representations in scene-selective occipital place area (OPA) are more highly correlated with representations from a network trained to predict navigational affordances. However, this study was limited to pre-defined regions of interest, while the task representations we identify span the entire brain. Consequently, the brain regions predicted by each model provide an atlas of neural representation of visual tasks and allow us to further study the representational relationships among tasks.

Independently of the brain, visual tasks have relationships among them. Task representations that are learned specifically for one task can be transferred to other tasks. Computer vision researchers commonly use transfer learning between tasks to save supervision and computational resources. In this vein, Zamir et al.^[18]^ recently showed that by standardizing model structure and measuring performance in transfer learning, one can generate a taxonomic map for task transfer learning (“Taskonomy”). This map provides an account of how much information is shared across different vision tasks. Given this global task structure, we can infer clusters of information defined by segregation of tasks, and then ask: does the brain represent visual information in the same task-relevant manner?

We compared the relationships between tasks using both brain representations and task learning. These comparisons reveal clustering of 2D tasks, 3D tasks, and semantic tasks. Compared to general encoding models, building individual encoding models and exploiting existing relationship among models has the potential to provide more in-depth understanding of the neural representation of visual information.

## 2 Methods

### 2.1 Encoding Model

To explore how and where visual features are represented in human scene processing, we extracted different features spaces describing each of the stimulus images and used them in an encoding model to predict brain responses. Our reasoning is as follows. If a feature is a good predictor of a specific brain region, information about that feature is likely encoded in that region. In this study, we first parameterized each image in the training set into values along different feature dimensions in a feature space. For example, if the feature space of interest is an intermediate layer in a task-driven network, we simply fed the image into the network and extracted its layer activation. These values are used as regressors in a ridge regression model to predict brain responses to that image. Performance from the validation data is used to choose the regularization parameter in the ridge regression model. We chose to use a ridge regression model instead of more complicated models in order to retain the interpretability of model weights, which may provide insights into the underlying dimensions of the brain responses. Each voxel’s regularization parameter was chosen independently via 7-fold cross-validation based on the prediction performance of the validation data. Model performance was evaluated on the test data using both Pearson’s correlation and coefficient of determination (*R*^2^). To determine the significance of the predictions, we calculated the *p*-values of each correlation coefficient and report false discovery rate (FDR) corrected *p*-values.

### 2.2 Feature Spaces

To simultaneously test representations from multiple 2D, and 3D vision tasks, we used the latent space features from each of the 19 tasks in Taskonomy^[18]^. The 19 tasks are: autoencoding, colorization, curvature estimation, denoising, depth estimation, edge detection (2D), edge detection (3D) or occlusion edges detection, keypoint detection (2D), keypoint detection (3D), depth, reshading, room layout estimation, segmentation (2D), segmentation (2.5D), surface normal estimation, vanishing point estimation, semantic segmentation, object classification and scene classification. In the Taskonomy training scheme, an intermediate latent space with fixed dimension (16 × 16 × 8) was enforced for each of these networks. We obtained these latent space activations by feeding our images into each pre-trained task-specific networks in the task bank provided with the Taskonomy paper. We selected 19 of the 25 tasks due to the concern that the remaining tasks might not relate to human vision. Examples of these less relevant tasks include jigsaw puzzle and camera pose estimation. We then built individual ridge regression models with the extracted latent features to predict brain responses and measured the correlation between the prediction and the true response in the held-out dataset.

### 2.3 Neural Data

The images used in this study are from a publicly available large-scale fMRI dataset, BOLD5000^[19]^. In the BOLD5000 study, participants’ brains were scanned while they presented with real-world images while they fixated at the center of the screen and judged how much they liked the image using a button press. Stimulus images in the BOLD5000 dataset were chosen from standard computer vision datasets (ImageNet, COCO and SUN). Within BOLD5000, we used the data from three participants viewing 4916 unique images. These 4916 image trials are separated into training, validation and testing sets randomly in each model fitting process. Region of interest (ROI) boundaries that identify category-selective brain regions in the whole-brain map presented in our results were generated directly from the ROI masks provided with the BOLD5000 dataset.

### 2.4 Task Similarity Computation

For each task, we took prediction performance scores across all voxels (dim 55,000). We set the score of a voxel to zero if the *p*-value of the correlation is above significance threshold (*p*<0.05, FDR corrected). This gave us a performance matrix of meaningful correlations of size *m* × *v*, where *m* is the number of tasks of interest and *v* is the number of voxels. To analyze the relationship between tasks based on neural representations, we computed pairwise similarity across tasks in the performance matrix using cosine distance. We also experimented with other distance or similarity functions but they did not show substantial differences. These pairwise similarities were then used to construct graphs and similarity trees among tasks.

## 3 Results

### 3.1 Model Prediction on ROIs

In Figure 1 we show the prediction accuracy measured using Pearson’s correlations coefficient. This was done for the 19 task-related feature spaces that were used to predict brain responses in predefined ROIs. Overall the predictions using these feature spaces—which come from mid-level computer vision tasks—show significant correlations with brain responses, except for the feature space from the curvature task. Among scene-selective regions, such as parahippocampal place area (PPA), retrosplenial complex (RSC), occipital place area (OPA), and lateral occipital complex (LOC), models with 3D features (e.g. Keypoints, Edges) show far better predictions than models with 2D features. This finding is consistent with the results of Lescroart and Gallant^[10]^. In contrast, within early visual areas the prediction results between 2D and 3D features are not differentiable. Across all ROIs, features from object and scene classification tasks provide the best predictions. For more scene specific tasks or semantic tasks such as 3D keypoints/edges, 2.5D/semantic segmentation, depth, distance, reshading, surface normal, room layout, vanishing points estimation, and object/scene classification, scene-selective regions are better predicted as compared to early visual areas. These patterns are consistent across all three participants. These results provide evidence that scene-selective areas show selectivity for scene-specific task representations. As such, models aiming to explain responses in these brain areas should minimally include representations from these scene-related visual tasks.

**Figure 1:**
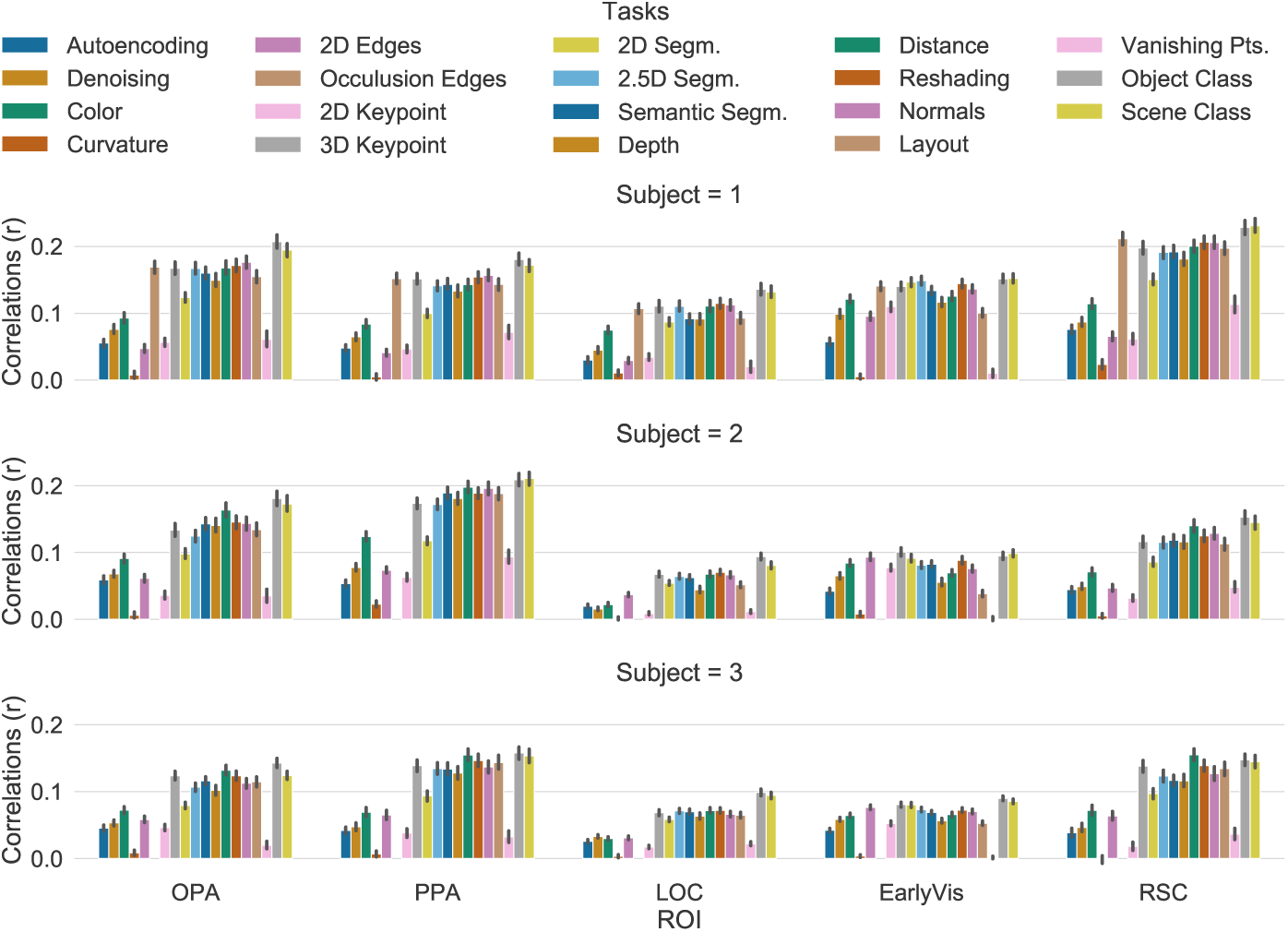
Pearson’s correlation coefficient of predicted response to true response across tasks. Three sub-figures correspond to prediction performance on three participants. Colors in the legend are arranged by columns. Features from 3D tasks, compared to those from 2D tasks, predict better in OPA, PPA, RSC, and LOC.

### 3.2 Model Prediction Across the Whole Brain

Prediction performance in pre-defined ROIs may omit relevant information arising in other brain regions. In Figure 2 and 3 we show prediction performance across the entire brain in a flattened view (generated using Pycortex^[20]^). Figure 2 shows the raw prediction performance as correlation coefficients for each task feature space. Figure 3 shows a contrast in prediction between 3D and 2D keypoints as well as edges. In this figure, red-colored voxels indicate that 3D features predict better than 2D features, while blue-colored voxels indicate that 2D features predict better than 3D features; white-colored voxels were predicted well by both features. We find that 3D features make better predictions for scene-selective regions—those delimited by ROI borders, while 3D and 2D features seem to predict early visual areas equally well. Both Figure 2 and 3 show that prediction results are consistent across three participants or tasks despite anatomical differences in their brain structures.

**Figure 2:**
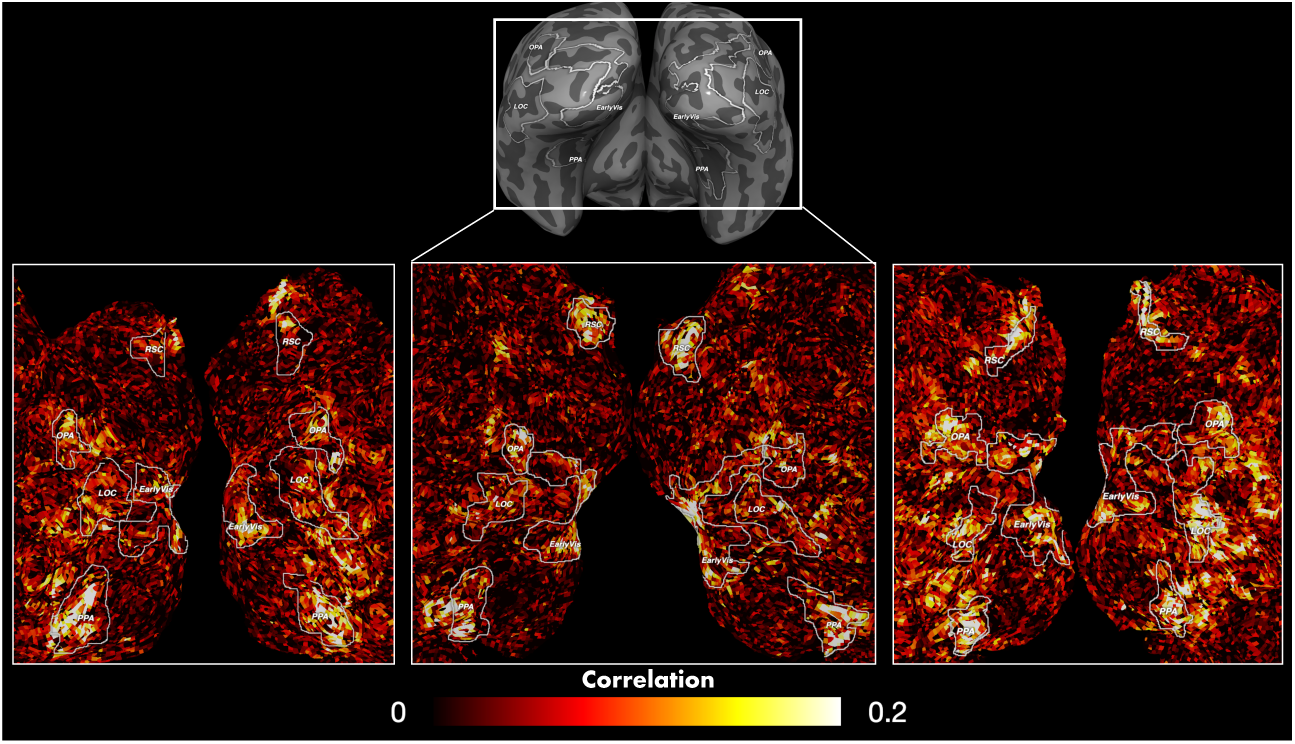
Whole brain prediction correlation using task presentation of scene classification network. The flat maps are cropped from the occipital regions of the brain. The upper zoom-out view shows the relative locations of the flat maps. Lower colored figures are the prediction performance across 3 participants. Prediction results are consistent across subjects.

**Figure 3:**
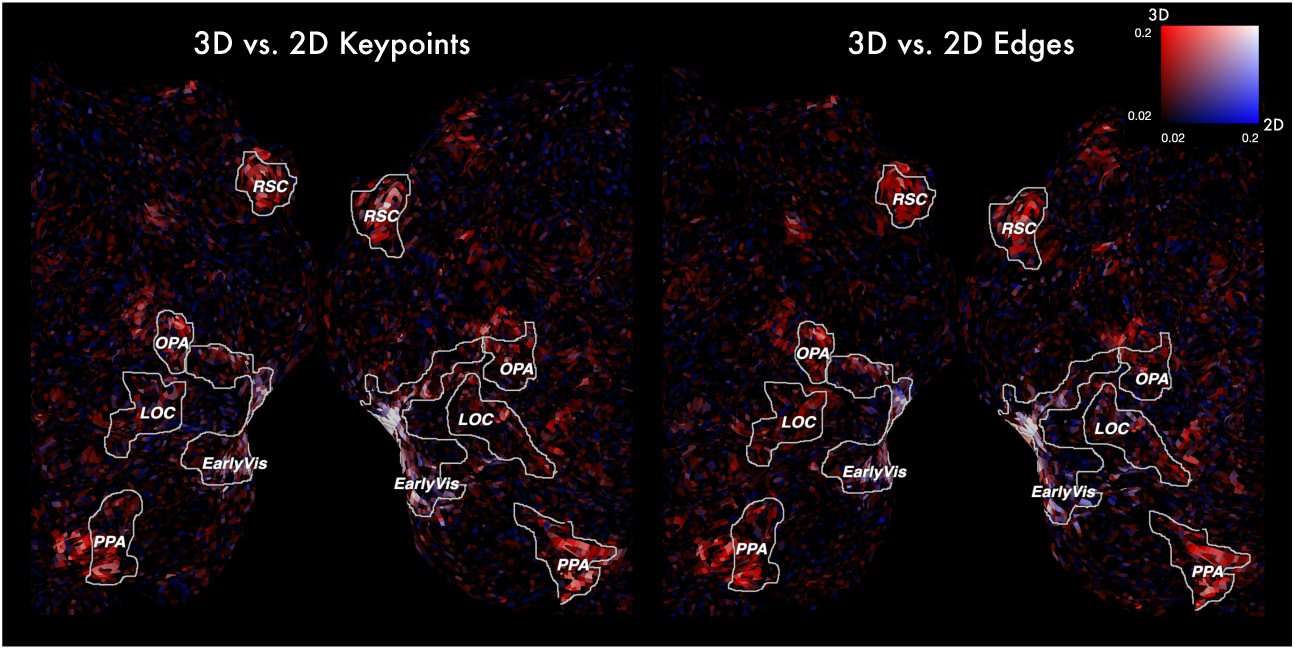
Contrast of prediction performance (meansured in Pearson’s correlation coefficients) between 2D and 3D features. The flat maps are cropped similarly as in Figure 2. 2D color map indicates the difference in correlation coefficients: red - 3D *>* 2D; blue - 2D *>* 3D. 3D task features predict better in scene selective regions and in more anterior parts of the brain.

Model performance using feature spaces from other tasks are shown in Figure 4. Here we plot 6 of the 19 tasks, while the remainder of the figures are provided in the appendix. Voxels with insignificant predictions (*p* > 0.05, FDR corrected) are not shown.

**Figure 4:**
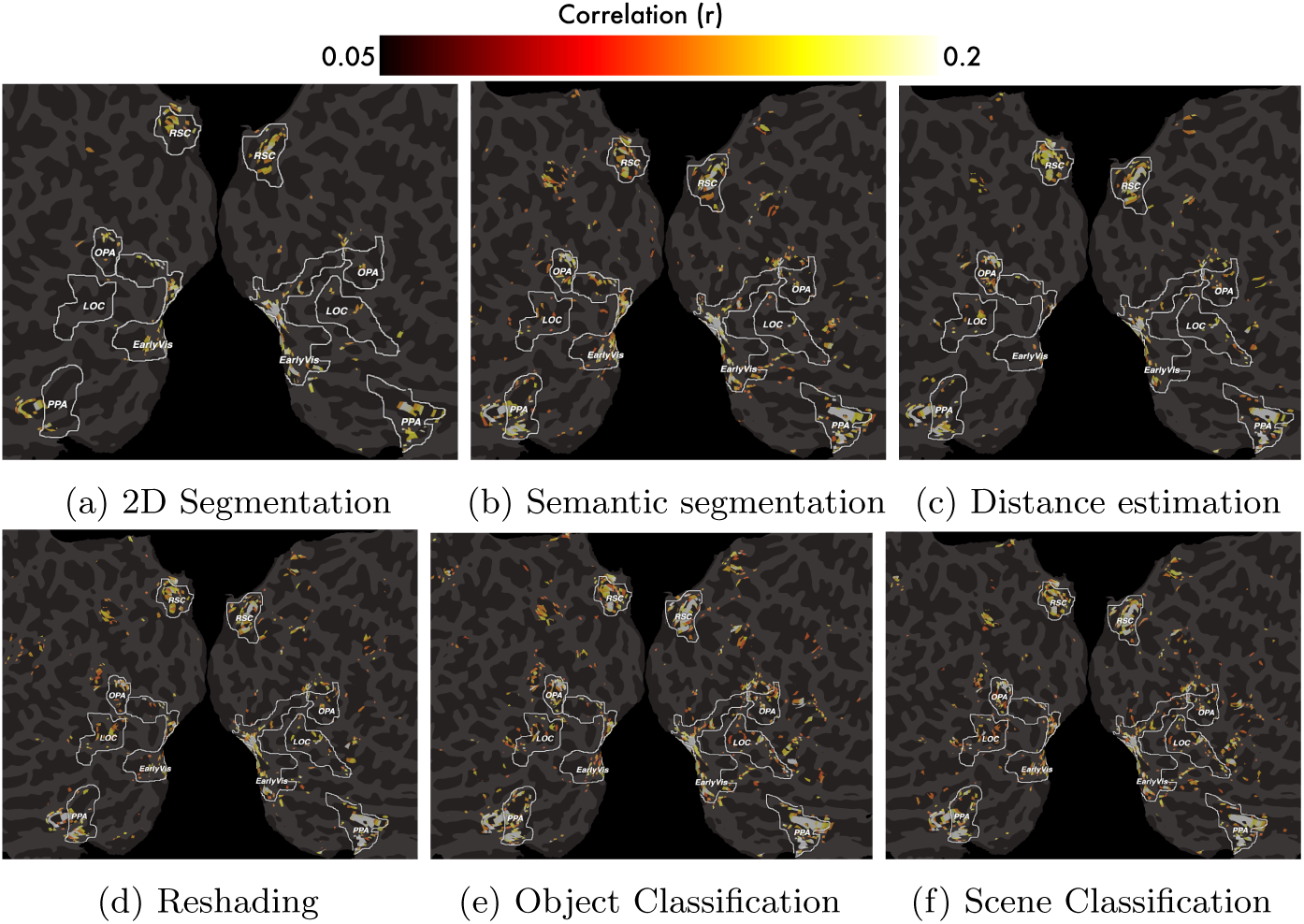
Predictive voxels using tasks features from Taxonomy^[18]^. Predictive regions of different tasks differ from each other but differences are not obvious.

### 3.3 Evaluation of Neural Representation Similarity

To this point we have shown that the neural prediction maps across tasks differ from one another; at the same time, there are many overlapping voxels across the predicted regions. Importantly, this pattern of voxels as predicted by the tasks can be exploited and used to infer task relationships in the brain. We computed task similarity using the methods discussed in 2.4 (Figure 5).

**Figure 5:**
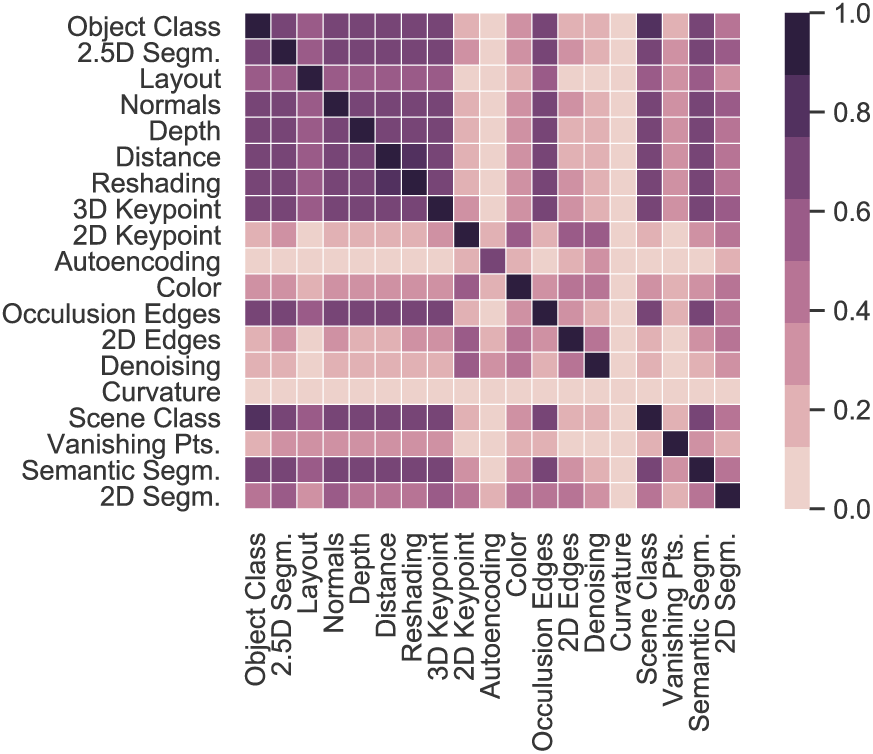
Task similarity matrix. Higher values means more similar.

### 3.4 Task Similarity Tree

To further explore the relationship between tasks as represented in the brain, we visualized the task similarity matrix using a dendrogram. Figure 6b shows the task similarity tree generated based on similarity in voxel prediction performance. Here, we compare these results with the task similarity tree based on transferring-out patterns in the original Taskonomy paper^[18]^, shown in Figure 6a. In the Taskonomy result, tasks are clustered into 3D, 2D, low-dimensional geometric and semantic tasks. Interestingly, the tree derived from brain representation also shows a similar structure: Semantic, 2D and 3D tasks were clustered together. The differences between two similarity trees may be due to low absolute model performance in the encoding model. For example, the model with feature from curvature estimation task has less than 10 significant voxels which may lead to some bias in the representation of the task tree. Overall the similarity between two task trees shows that, at a coarse level, neural representation of task information is similar to that found through transfer learning.

**Figure 6:**
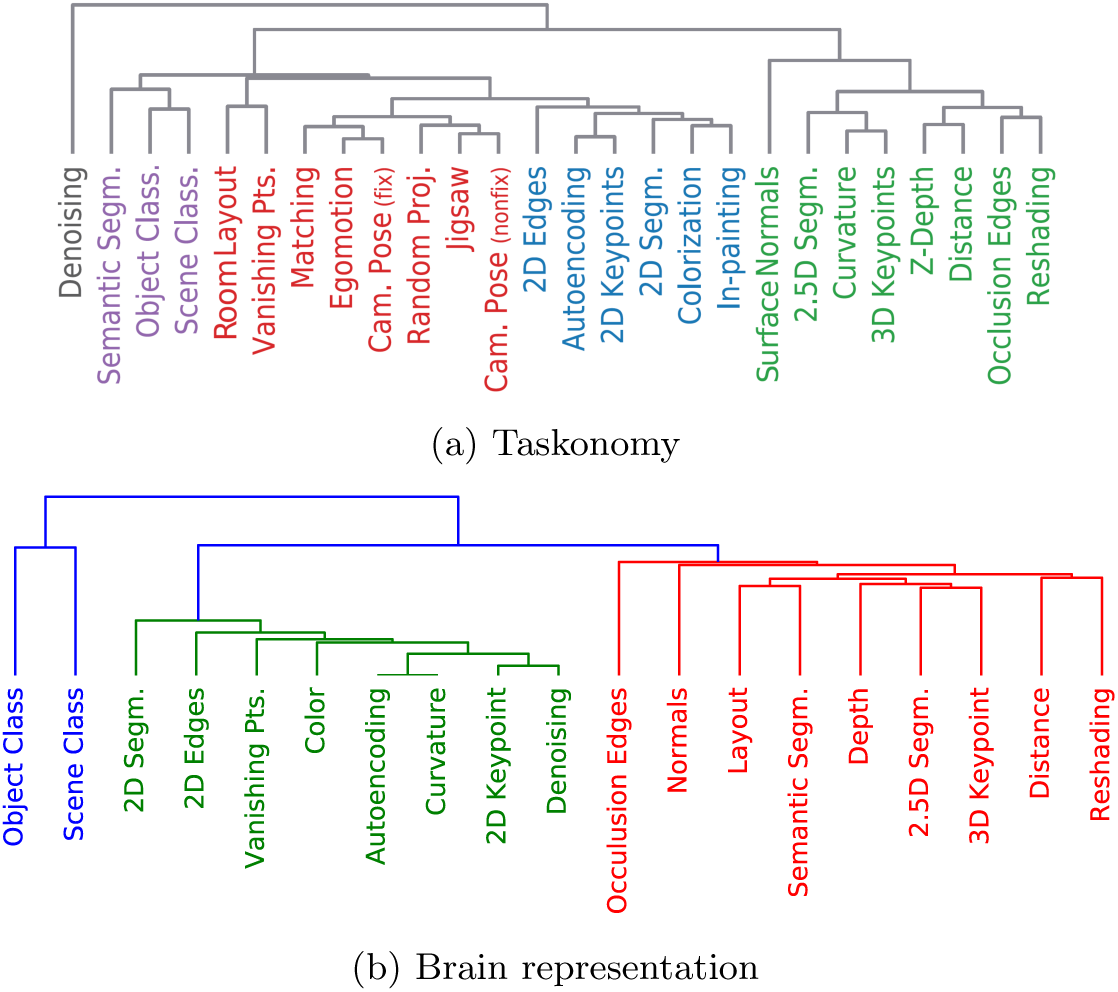
Task Trees from Taxonomy^[18]^ and brain representation. Similar clusters of 2D, 3D and semantic tasks are found in both trees.

### 3.5 Task Graphs

In order to explore the similarity pattern across tasks at a more fine-grained level, we compared the Taskonomy graph (Figure 7a) to its brain-derived version (Figure 7b). In the brain-derived task graph, widths of edges between two nodes are measured by the cosine similarity in voxel prediction performance as described in 2.4. The graph shown here for Participant 1 is consistent with those of the other two participants, indicating that the graphs capture a meaningful representational pattern in brain responses. Interestingly, similar edges exist between the Taskonomy and brain graphs. For example, heavy edges connect between occlusion edges and surface normal, reshading, and 3D keypoint, *et. cetera*.

**Figure 7:**
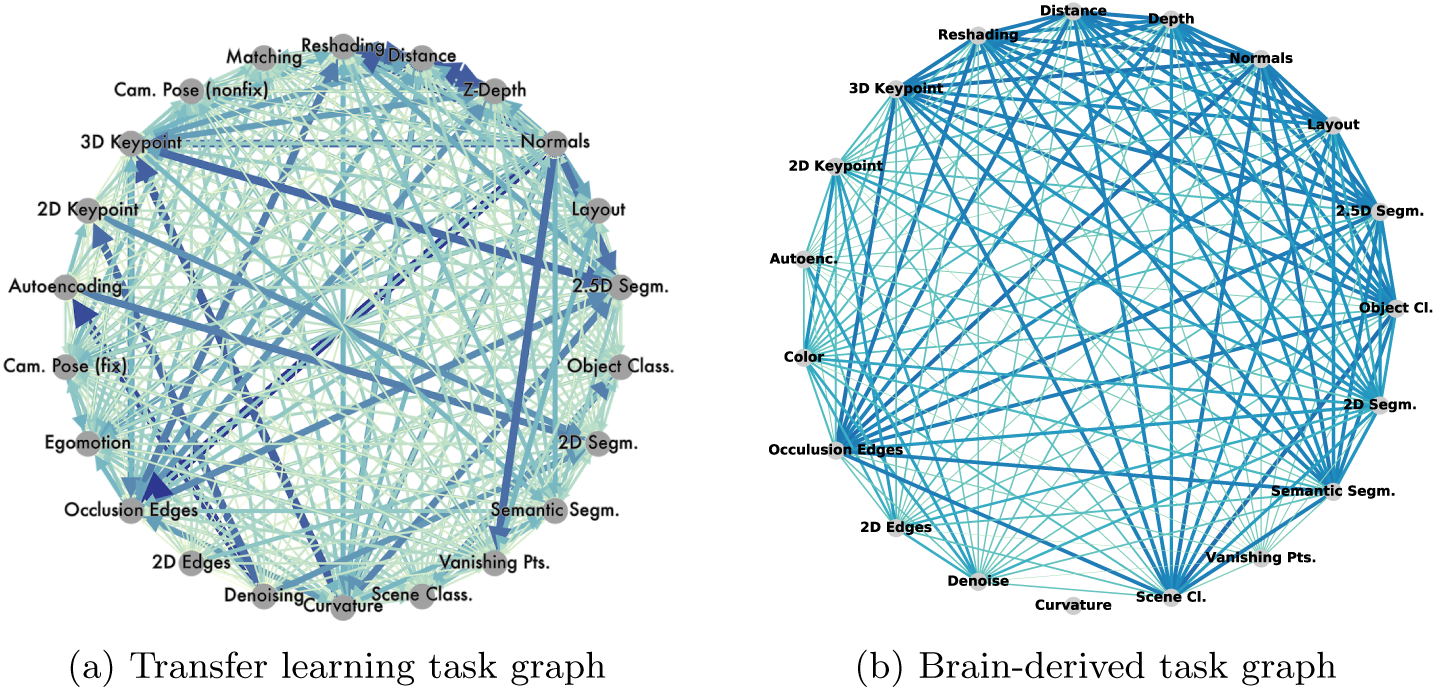
Task networks. (a) Task graph from Taskonomy^[18]^. (b) Task graph computed using similarity in predictive brain regions across tasks. Widths of edges between two nodes are measured by the cosine similarity in voxel prediction performance as described in 2.4

## 4 Discussion

There is growing evidence that the architecture of the primate visual system reflects a series of computational mechanisms that enable high performance for accomplishing evolutionarily adaptive tasks^[21]^. However, the precise nature of these tasks remains challenging problem for many reasons, including the limitations of neuroscience data collection methods and the lack of interpretability of intermediate representations in both biological and artificial systems. To address these challenges we leveraged both the space of vision tasks learned through transfer learning in Taskonomy^[18]^ and the recent availability of one of the first larger-scale human functional neural datasets, BOLD5000^[19]^. However, there are substantial differences in the image distribution of BOLD5000, which contains general objects and scenes, and the Taskonomy dataset, which includes exclusively indoor scenes. Note that when we applied the pre-trained Taskonomy models to BOLD5000 images, we found that these models performed more poorly as compared to when using the original Taskonomy dataset, especially for the outdoor images in BOLD5000. Such inconsistency in image distribution is unavoidably reflected in the encoding model performance and hinders us from making more specific claims about task spaces in the brain. One solution to this issue would be to use a more general computational model of visual tasks, as well as a larger brain dataset based on more images, both of which are outside of the scope of this paper.

## 5 Conclusion

Our results reveal that task-specific representations in neural networks are useful in predicting brain responses and localizing task-related information in the brain. One of the main findings is that features from 3D tasks, compared to those from 2D tasks, predict a distinct part of visual cortex. We also explored the fine-grained neural representation of all 19 Taskonomy tasks with similarity trees and graphs. In the future we will incorporate features from other tasks to obtain a more comprehensive picture of task representation in the brain.

For years neuroscientists haven been focused on recovering which parts of the brain do what. However, what are the computational principles behind the encoding of information in the brain? We observe feedforward hierarchies in the visual pathways, but what are the stages of information processing? To date, we have few satisfying answers. The ultimate goal in studying task representation in the brain is to answer some of these questions. We exploited the task relationship found in transfer learning and used it as a ground truth of visual information space to understand the neural representation of visual and semantic information. In sum, our paper provides a first attempt in using predictive-map-generated task relationships to answer broader questions of neural information processing.

## References

[1] Russell Epstein and Nancy Kanwisher. A cortical representation of the local visual environment. Nature, 392(6676):598, 1998.

[2] Dwight J Kravitz, Cynthia S Peng, and Chris I Baker. Real-world scene representations in high-level visual cortex: it’s the spaces more than the places. Journal of Neuroscience, 31(20):7322–7333, 2011.

[3] Soojin Park, Talia Konkle, and Aude Oliva. Parametric coding of the size and clutter of natural scenes in the human brain. Cerebral cortex, 25(7): 1792–1805, 2014.

[4] Assaf Harel, Dwight J Kravitz, and Chris I Baker. Deconstructing visual scenes in cortex: gradients of object and spatial layout information. Cerebral Cortex, 23(4):947–957, 2012.

[5] Mark D Lescroart, Dustin E Stansbury, and Jack L Gallant. Fourier power, subjective distance, and object categories all provide plausible models of bold responses in scene-selective visual areas. Frontiers in computational neuroscience, 9:135, 2015.

[6] Katrina Ferrara and Soojin Park. Neural representation of scene boundaries. Neuropsychologia, 89:180–190, 2016.

[7] Frederik S Kamps, Joshua B Julian, Jonas Kubilius, Nancy Kanwisher, and Daniel D Dilks. The occipital place area represents the local elements of scenes. Neuroimage, 132:417–424, 2016.

[8] Simon Kornblith, Xueqi Cheng, Shay Ohayon, and Doris Y Tsao. A network for scene processing in the macaque temporal lobe. Neuron, 79(4):766–781, 2013.

[9] Michael F Bonner and Russell A Epstein. Coding of navigational affordances in the human visual system. Proceedings of the National Academy of Sciences, 114(18):4793–4798, 2017.

[10] Mark D Lescroart and Jack L Gallant. Human scene-selective areas represent 3d configurations of surfaces. Neuron, 101(1):178–192, 2019.

[11] Dustin E Stansbury, Thomas Naselaris, and Jack L Gallant. Natural scene statistics account for the representation of scene categories in human visual cortex. Neuron, 79(5):1025–1034, 2013.

[12] Pulkit Agrawal, Dustin Stansbury, Jitendra Malik, and Jack L Gallant. Pixels to voxels: Modeling visual representation in the human brain. arXiv preprint 1407.5104, 2014.

[13] Daniel LK Yamins, Ha Hong, Charles F Cadieu, Ethan A Solomon, Darren Seibert, and James J DiCarlo. Performance-optimized hierarchical models predict neural responses in higher visual cortex. Proceedings of the National Academy of Sciences, 111(23):8619–8624, 2014.

[14] Umut Güçlü and Marcel AJ van Gerven. Deep neural networks reveal a gradient in the complexity of neural representations across the ventral stream. Journal of Neuroscience, 35(27):10005–10014, 2015.

[15] Michael Eickenberg, Alexandre Gramfort, Gaël Varoquaux, and Bertrand Thirion. Seeing it all: Convolutional network layers map the function of the human visual system. NeuroImage, 152:184–194, 2017.

[16] Alexander JE Kell, Daniel LK Yamins, Erica N Shook, Sam V Norman-Haignere, and Josh H McDermott. A task-optimized neural network replicates human auditory behavior, predicts brain responses, and reveals a cortical processing hierarchy. Neuron, 98(3):630–644, 2018.

[17] Kshitij Dwivedi and Gemma Roig. Task-specific vision models explain task-specific areas of visual cortex. BioRxiv, page 402735, 2018.

[18] Amir R Zamir, Alexander Sax, William Shen, Leonidas J Guibas, Jitendra Malik, and Silvio Savarese. Taskonomy: Disentangling task transfer learning. In Proceedings of the IEEE Conference on Computer Vision and Pattern Recognition, pages 3712–3722, 2018.

[19] Nadine Chang, John A Pyles, Austin Marcus, Abhinav Gupta, Michael J Tarr, and Elissa M Aminoff. Bold5000, a public fmri dataset while viewing 5000 visual images. Scientific data, 6(1):49, 2019.

[20] James S Gao, Alexander G Huth, Mark D Lescroart, and Jack L Gallant. Pycortex: an interactive surface visualizer for fmri. Frontiers in neuroinformatics, 9:23, 2015.

[21] Daniel LK Yamins and James J DiCarlo. Using goal-driven deep learning models to understand sensory cortex. Nature neuroscience, 19(3):356, 2016.

