## Appendix for "Neural Taskonomy: Inferring the Similarity of Task-Derived Representations from Brain Activity"

### 1 Appendix

#### 1.1 Model predictions of tasks from Taskonomy

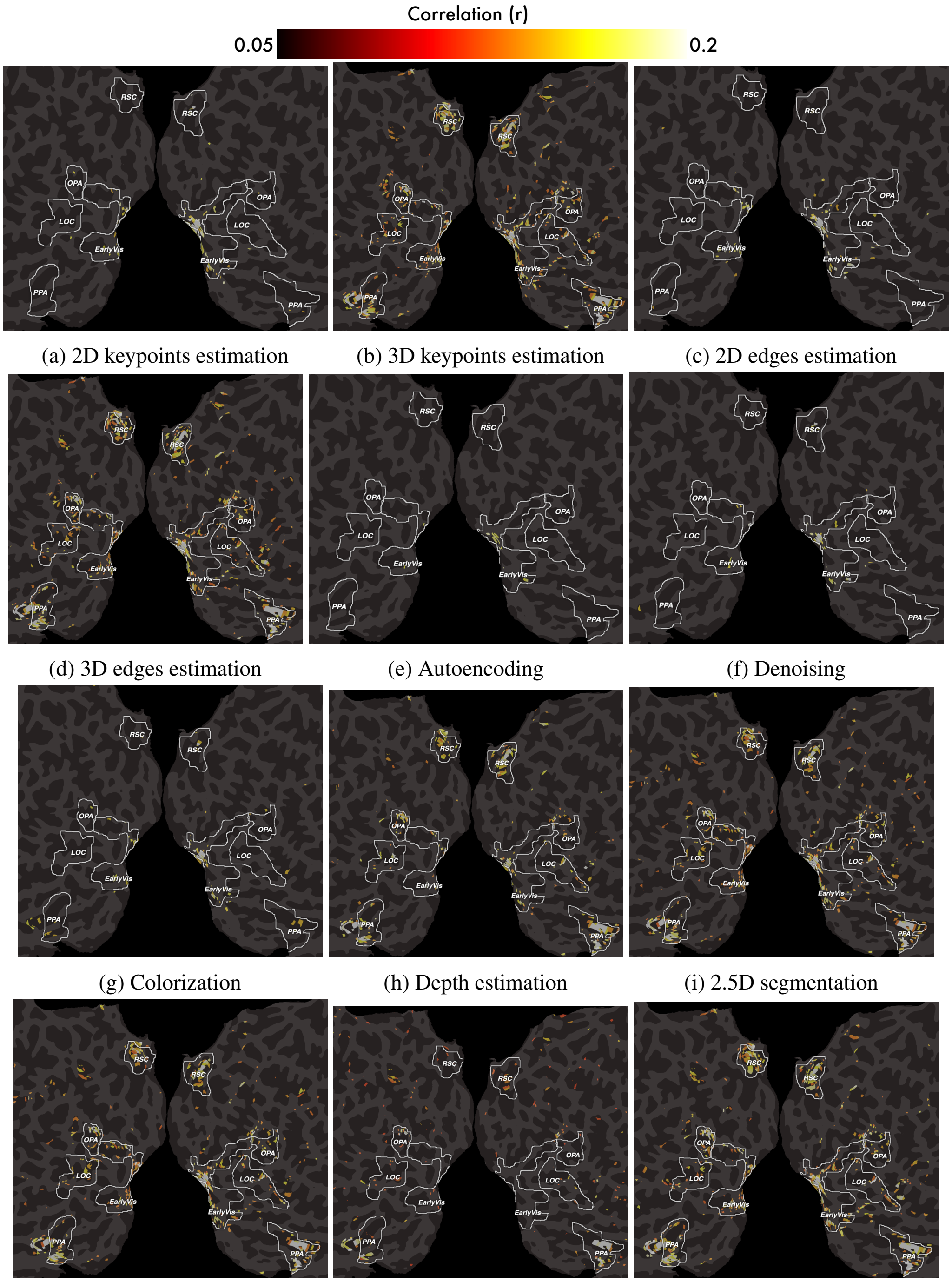


(a) 2D keypoints estimation (b) 3D keypoints estimation (c) 2D edges estimation

(d) 3D edges estimation (e) Autoencoding (f) Denoising

(g) Colorization (h) Depth estimation (i) 2.5D segmentation

(j) 2.5D segmentation (k) Vanishing point estimation (l) Surface normal estimation

Figure 1: Model predictions of tasks from Taskonomy. Only voxels above significance threshold ($p<0.05$, FDR corrected) are shown here.
